## Supplemental data for "Designing efficient genetic code expansion in Bacillus subtilis to gain biological insights"

### Supplemental Text

##### Promoter Screen

Five different combinations of AARS and tRNA promoter were constructed and tested for activity, as it has been previously demonstrated that optimization of expression levels is necessary for good activity[1]. These combinations were tested and a pVeg constitutive promoter in front of the synthetase and a pSer promoter in front of the tRNA were found to be the most effective (Supp. Fig. 1A).

##### Reporter optimization

Initial experiments with the mNeongreen reporter expressed by the IPTG-inducible pHyperspank showed high levels of background in the absence of nsAA & synthetase. Follow-up experiments indicated that the background was due to a secondary start codon at Methionine10 of mNeongreen, driven by a ribosomal binding site ~1/3 of the strength of the canonical pHyperspank contained in residues 4-8 (Supp. Fig. 1B-C). Ribosomal binding site presence and strength was calculated from the Salis lab ribosomal binding site calculator[2]. These findings could drive reinterpretation of subcellular microscopy experiments that used C-terminal or sandwich fusion mNeongreen tags[3,4], as the commonly used mNeongreen sequence is capable of initiating translation independently. An M10S mutation suppressed the background and reported 30-50 fold increase in mNeongreen fluorescence upon addition of the nsAA (Supp. Fig. 1C). All subsequent usage of mNeongreen used the M10S variant.

##### Synthetase Promiscuity

Further exploration of general nsAA incorporation in *B. subtilis* with the extremely sensitive nanoluciferase reporter[5] showed subtleties of incorporation efficiency and promiscuous background incorporation. The MjTyrRSs are known to promiscuously incorporate native amino acids in the absence of the target nsAAs[6], which is reflected here by some background incorporation in the absence of nsAA. When tested with the UAG-nanoluciferase reporter, the presence of napARS increased UAG-luciferase expression 22-fold in the absence of nsAA. The addition of nsAA increased expression another 25-fold, for a total of 556-fold increase over the UAG-nanoluciferase reporter alone. Both the ScwRS and MaPylRS showed minimal background incorporation, with addition of the synthetase causing 1.3 and 3.6-fold over the reporter in the absence of nsAA, respectively. Addition of nsAA increased expression 11 and 295-fold over the reporter in the presence of corresponding nsAAs, respectively (Supp. Figure 1G).

### Supplemental Movies

#### Supplemental Movie 1

Titration of MciZ expression and its effects on FtsZ filaments in vivo. Cells expressing mNeonGreen-FtsZ were imaged at 1-second intervals for 100 seconds by TIRF microscopy. The concentration of pAzF added is indicated in each panel. In each case, the UAG-MciZ construct was induced with 100 µM IPTG. The movie is displayed at 30 frames per second (30x actual speed). Scale bar: 2 µm.

### Supplemental Figures

#### Supplemental Figure 1

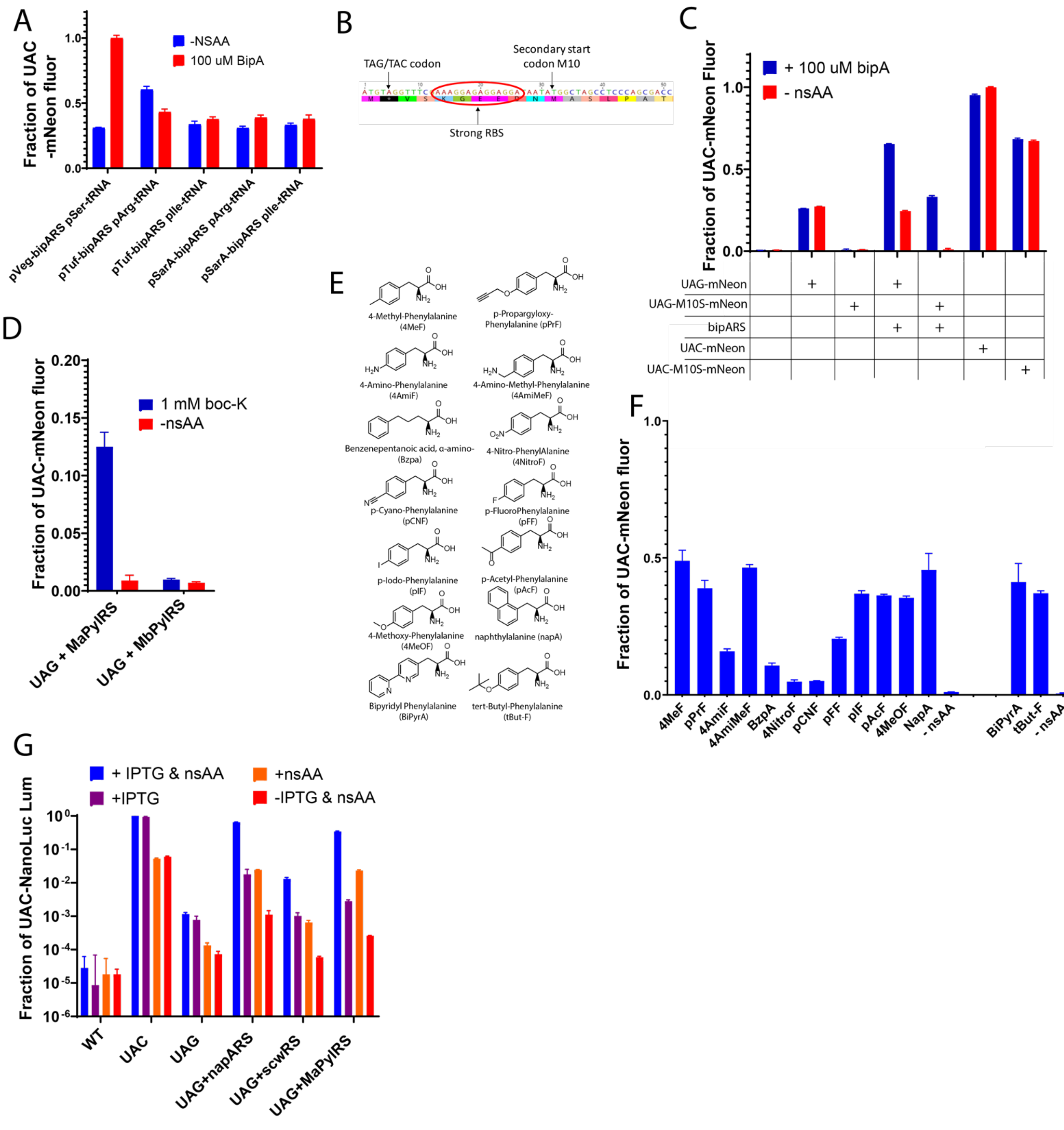


**Supplemental Figure 1:** ***Extended nsAA incorporation in B. subtilis*****.** For all bar graphs, average of three replicates is shown with standard deviation as error bar. Fluor indicates mNeongreen fluorescence, while Lum indicates nanoLuciferase luminescence. 100 uM of nsAA was used for all tyrosine-based nsAAs, but 1 mM was used for nsAAs **5** & **6** A). Assay of 5 AARS/tRNA promoter combinations, with the identity of the promoter indicated below the bars. Reported by an pHyperspank-inducible mNeongreen containing a UAG codon in an N-terminal linker and normalized to maximum fluorescence from the experiment. B) Schematic of first 17 residues of TAG-mNeongreen reporter, with the M10 capable of secondary translational start indicated. Secondary RBS strength was calculated as approximately 1/3 the strength of the pHyperspank RBS with the Salis Lab Ribosomal Biding site calculator. C) Assay of the mNeongreen and the M10S mNeongreen reporters with and without associated synthetase. The reported TAG or TAC codon is at position 2, after the start methionine. D) Comparison of MaPylRS and MbPylRS activity with the TAG-M10SmNeongreen reporter, normalized to TAC-M10SmNeongreen. E) Structures and names of additional nsAAs incorporated in F) using the napARS synthetase (bars on left) or the bipARS synthetase (bars on right). G) Assay of 3 different synthetases incorporating nsAA **1** for napARS, **5** for MaPylRS and **6** for ScwRS using a sensitive IPTG-inducible TAG-nanoluciferase reporter capable of reporting over 5 orders of magnitude. Log plot of pHyperspank-Nanoluciferase levels with a UAG and a UAC codon to demonstrate accurate levels of promiscuity vs. native amino acids.

#### Supplemental Figure 2

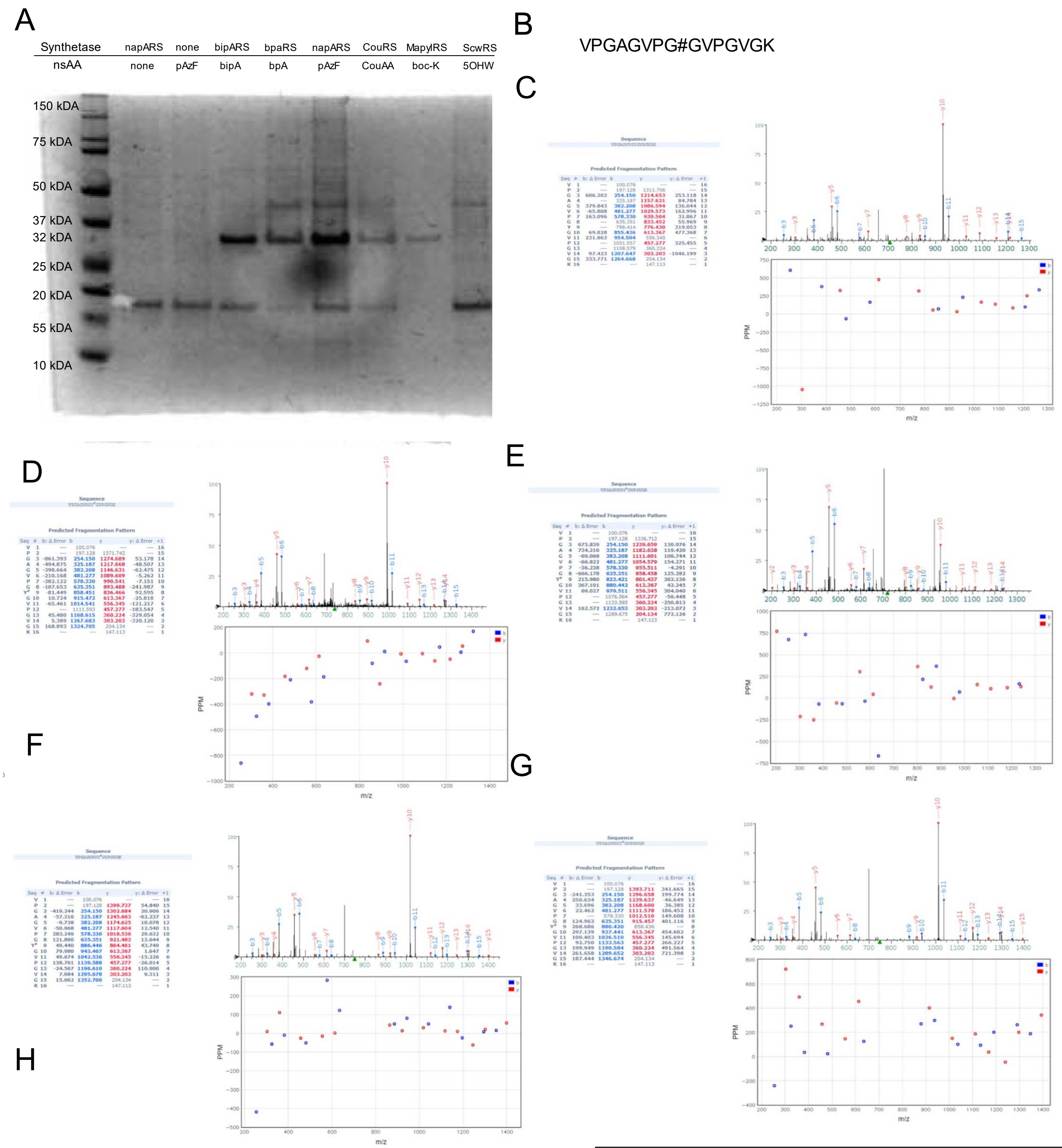


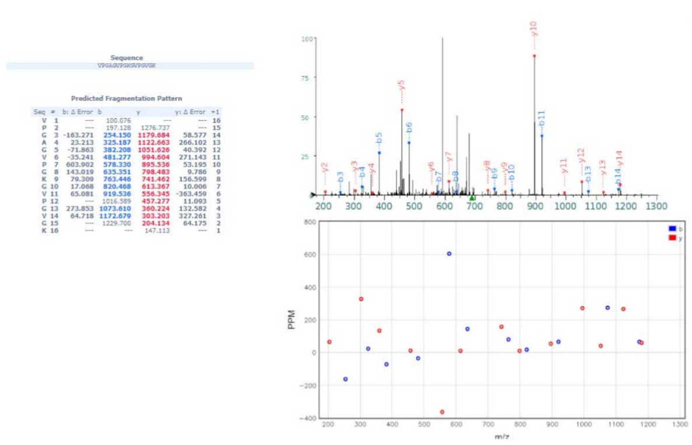

**Supplemental Figure 2: Mass-spectrometry of nsAA-containing peptides.** A) Gel image showing purified mNeongreen containing nsAAs. B) Sequence of 16-resitdue Elastin-like peptides found in mass-spec, with position of nsAA indicated by #.C-H) Mass Spectrums of peptides containing C) Tyrosine D) pAzF E) bipA F)pBpA G) CouAA H) Lysine (likely resulting from deprotection of boc-lysine)

**Supplemental Figure 3**

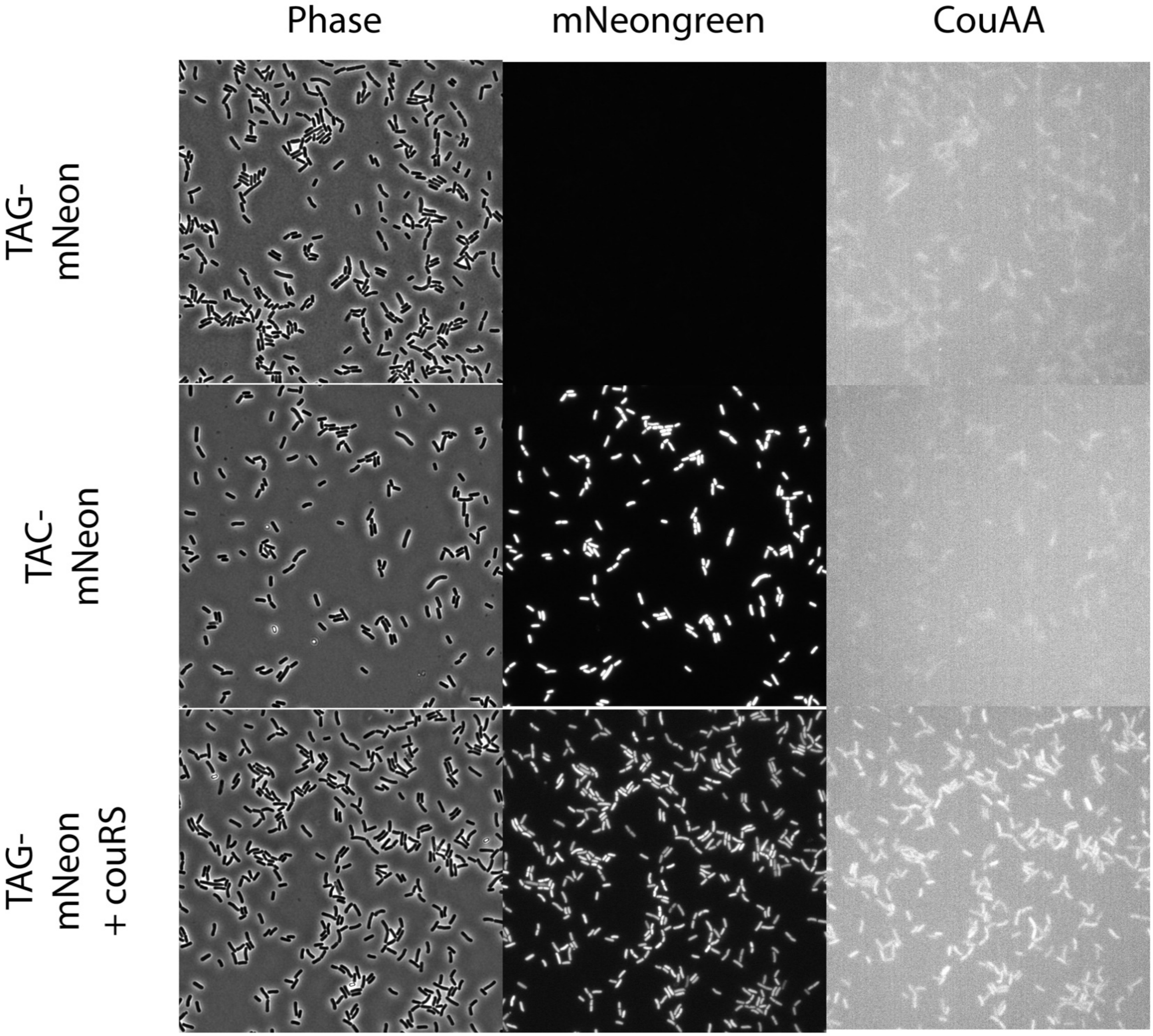


**Supplemental Figure 3: Fluorescence imaging using CouAA** Images of *B. subtilis* taken in phase, GFP (mNeongreen), and DAPI (CouAA) wavelengths. Each column has identical imaging conditions and brightness/contrast settings.

#### Supplemental Figure 4

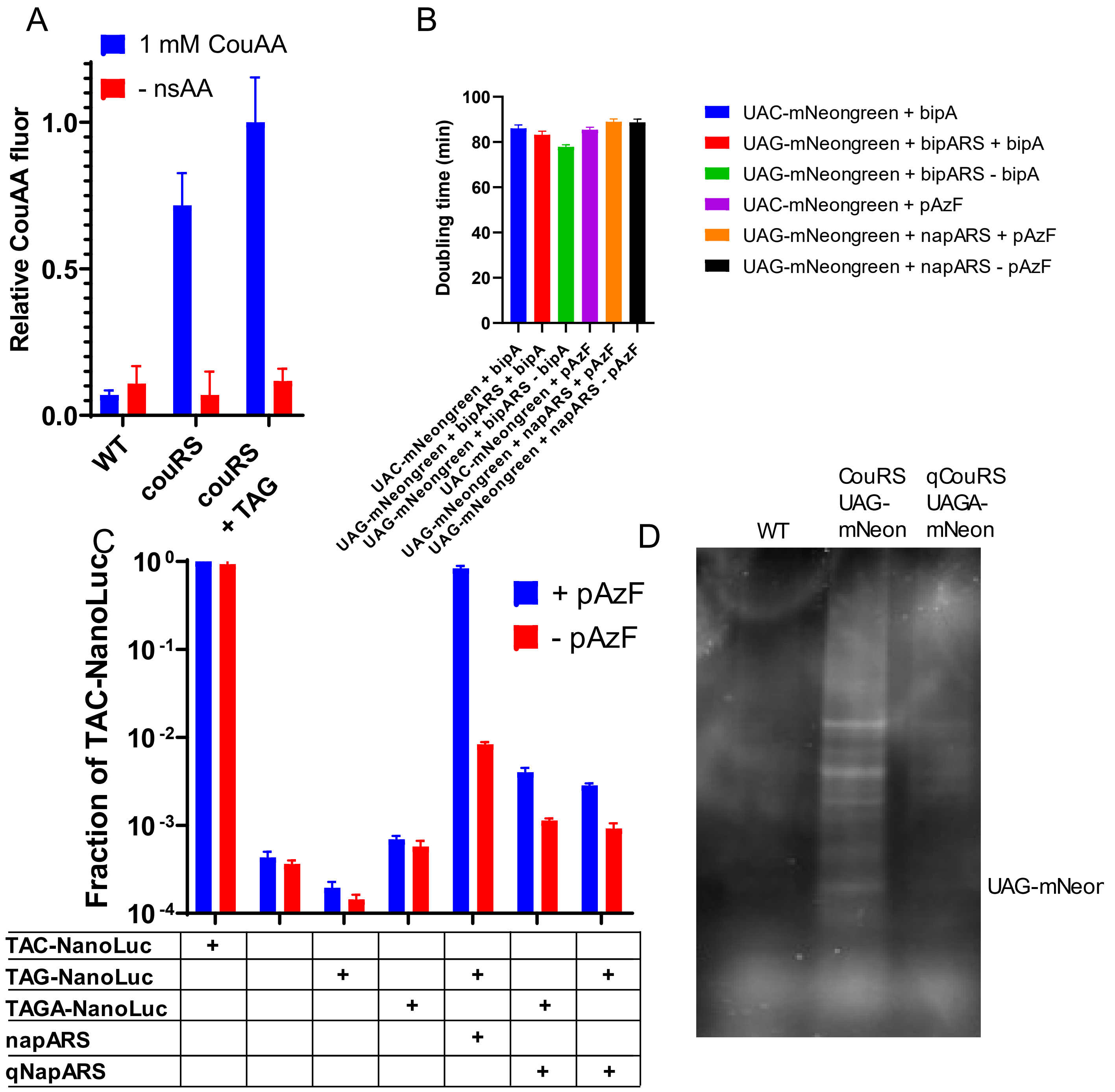


**Supplemental Figure 4: Genomic incorporation and doubling times.** A) Fluorescence of the CouAA amino acid remaining in bulk cells after washing. Average of triplicates shown with standard deviation as the error bar, normalized to fraction of UAG signal. B) Doubling times of *B. subtilis* cells with and without synthetase and nsAA. C) High-sensitivity detection of nanoluciferase reporter with UAC, UAG or UAGA codons inserted at position 2 of NanoLuc. Average of triplicates shown with standard deviation as the error bar. Normalized to fraction of expression of UAC construct. D) Whole-cell lysate of cells grown with CouAA run on SDS-page gels and imaged for CouAA fluorescence in the proteome.

#### Supplemental Table 1

| # | Gene Name | Percentage of nsAA-containing UAG proteins | Gene function | Essential? | Ends with TAGA? |
|---|---|---|---|---|---|
| 1 | alsS | 13.1 | acetolactate synthase | No | No |
| 2 | purB | 10.0 | purine biosynthesis | No | Yes |
| 3 | rplX | 5.1 | ribosomal protein L24 | Yes | No |
| 4 | rpsG | 4.5 | ribosomal protein S7 | Yes | No |
| 5 | yfmC | 4.0 | iron uptake | No | No |
| 6 | secA | 3.6 | protein secretion | No | No |
| 7 | accD | 3.4 | acetyl-CoA carboxylase | Yes | Yes |
| 8 | ytpR | 3.2 | unknown (possible tRNA synthetase) | No | Yes |
| 9 | atpG | 3.0 | ATP synthase | No | Yes |
| 10 | pyrAA | 3.0 | pyrimidine biosynthesis | No | Yes |
| 11 | ilvD | 2.9 | biosynthesis of branched-chain amino acids | No | Yes |
| 12 | yloV | 1.7 | fatty acid kinase | No | Yes |
| 13 | dapB | 1.6 | biosynthesis of lysine and peptidoglycan | Yes | Yes |
| 14 | ywlF | 1.5 | ribose-5-phosphate isomerase | No | No |
| 15 | ywhA | 1.4 | unknown (probable transcriptional regulator) | No | No |
| 16 | yaaK | 1.4 | unknown (possible DNA-binding protein) | No | No |
| 17 | murE | 1.3 | peptidoglycan precursor biosynthesis | Yes | No |
| 18 | ylxF | 1.2 | unknown | No | No |
| 19 | noc | 1.2 | control of cell division | No | Yes |
| 20 | ylxM | 1.1 | presecretory protein translocation | No | No |
| 21 | aroC | 1.1 | biosynthesis of aromatic amino acids | No | Yes |
| 22 | gsaB | 1.0 | biosynthesis of heme, modification of Efp | No | Yes |
| 23 | yeeB | 1.0 | unknown (possible helicase) | No | No |

**Supplemental Table 1: High-abundance proteins containing nsAAs.** Proteins containing UAG stop codons ranked by abundance in the nsAA-incorporation enrichment by click-pulldown and quantified by mass-spectrometry. Only proteins above 1% abundance are shown and make up 71% of all nsAA-containing proteins found in the enrichment.

#### Supplemental Figure 5

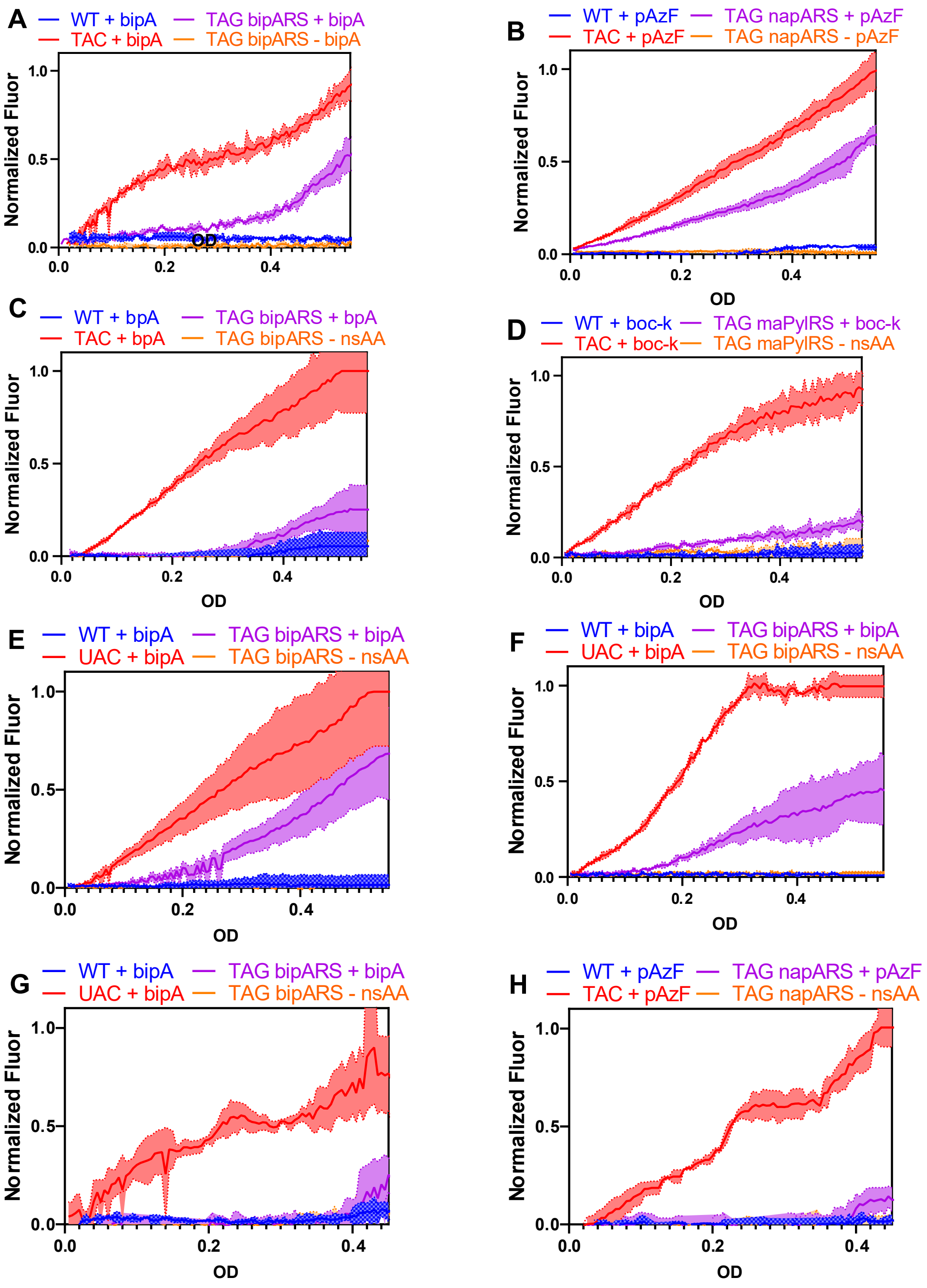


**Supplemental Figure 5: Fluorescence vs. OD time courses for various nsAAs.** Fluorescence and OD curves for different nsAAs and media conditions. A-D) S750 minimal media. E) S750 minimal media plus 1% w/w Pluronic F-68 F) S750 media modified to replace all amino acids with 0.3% buffered ammonium sulfate. G-H) CH rich media.

#### Supplemental Figure 6

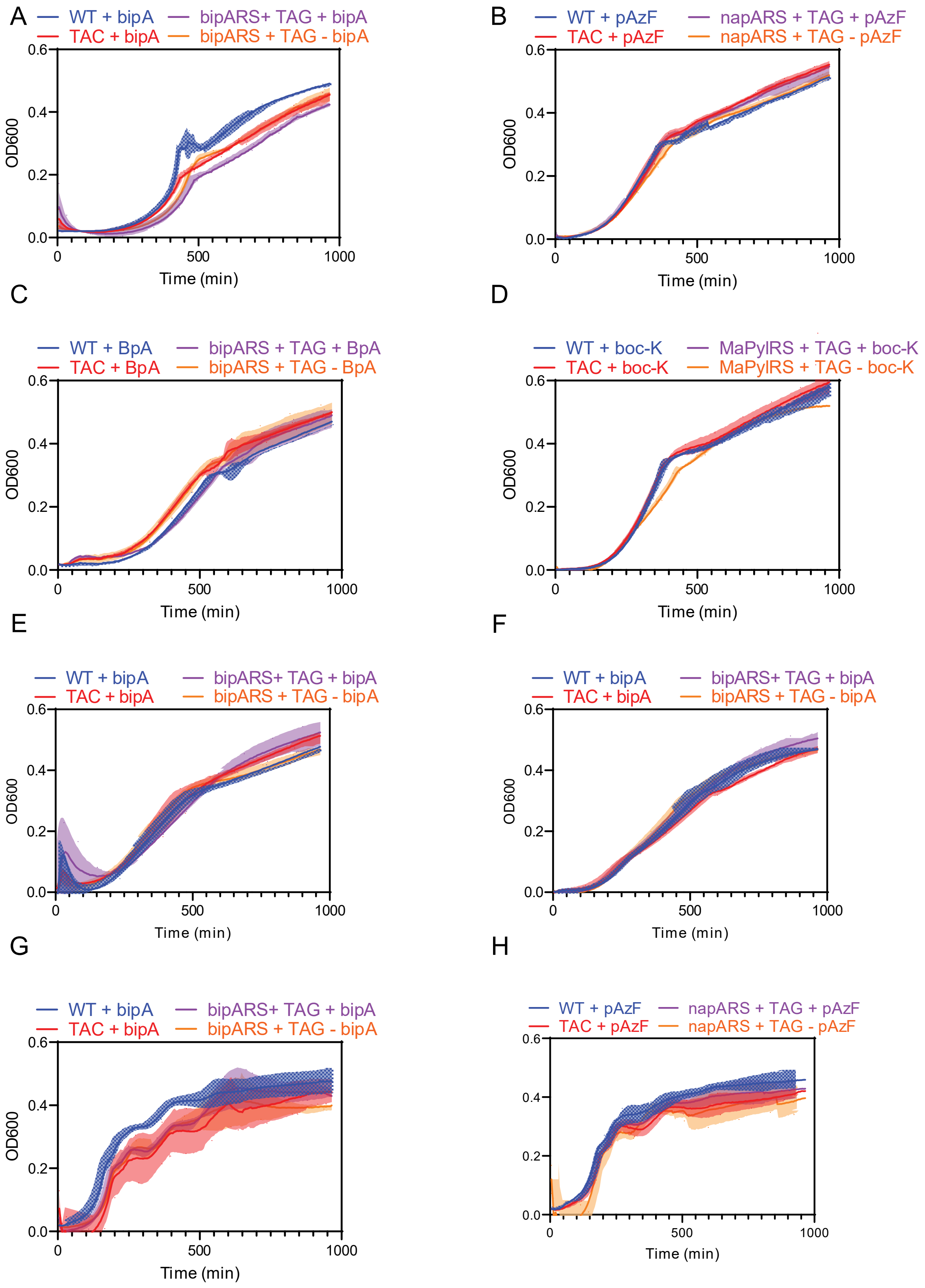


**Supplemental Figure 6. OD time courses for various nsAAs.** OD curves for different nsAAs and media conditions shown in Supplemental figure 5. A-D) S750 minimal media. E) S750 minimal media plus 1% w/w Pluronic F-68 F) S750 media modified to replace all amino acids with 0.3% ammonium sulfate. G-H) CH rich media.

**Supplemental Figure 7**

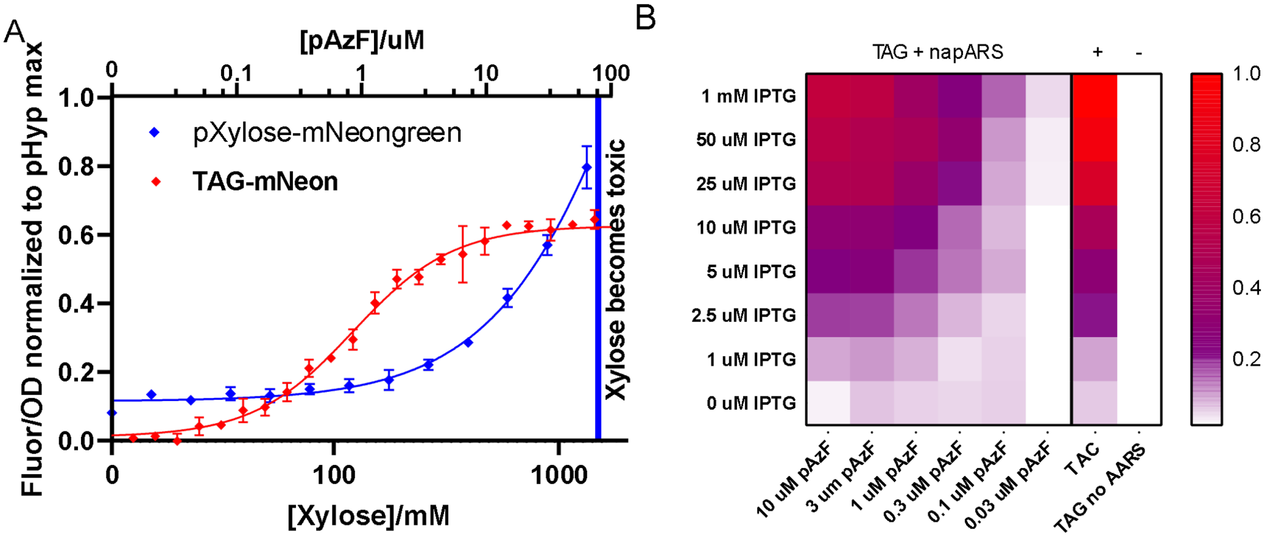


**Supplemental Figure 7**: Extended titration data. A) Titration and sigmoidal fit of pXylose-mNeongreen, with pAzF-induced TAG-mNeongreen from Figure 3A overlaid. 3M xylose significantly reduces growth rate B) 2-dimensional titration data, alternate display for the dataset in figure 3B.

**Supplemental Figure 8**

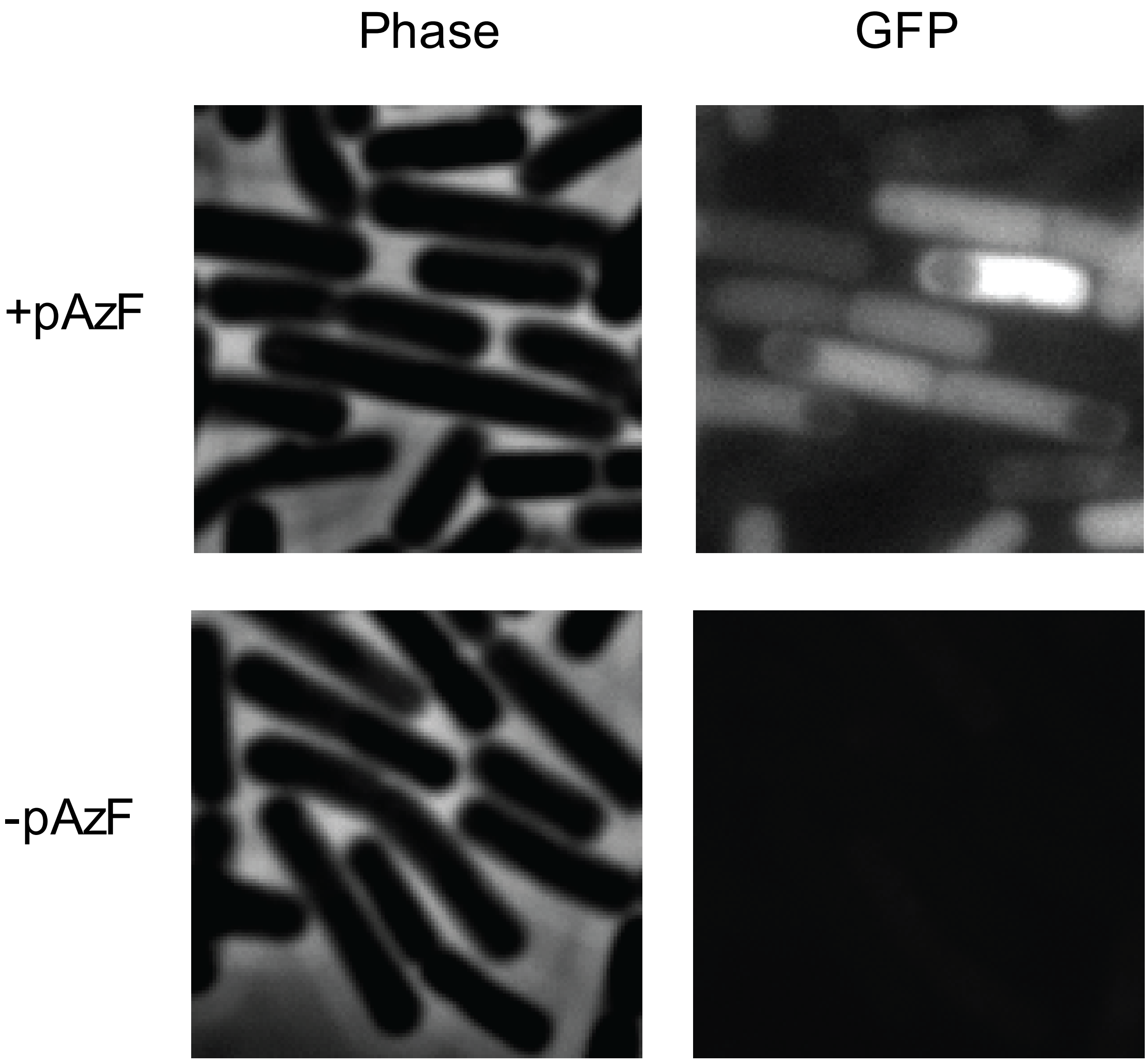


**Supplemental Figure 8: Incorporation of pAzF in sporulating *B. subtilis* cells.** Cells with GFP(F27TAG) under control of a mother-cell specific promoter, $P_{spoIIE}$, were induced to sporulate by resuspension. At the time of resuspension, the culture was split in two, and pAzF was added to the experimental sample. An example image of sporulating cells at 150 minutes after resuspension shows the GFP signal in the mother cell compartment of engulfed sporulating cells. GFP images shown were taken with identical acquisition times and settings. The control sample shows fluorescent signal at background levels when pAzF is not present.

#### Methods
